# Sundoli: A necro(w)bot with multi-stiffness joints built using geared mechanical metastructures

**DOI:** 10.1101/2025.05.16.654441

**Authors:** Hyeon Lee, Jonah Mack, Parvez Alam

## Abstract

In this communication, we design and analyse Sundoli, a necrobot (a bionically engineered robot using decreased animal parts). It is manufactured using a crow endoskeleton, supported and rearticulated by a geared mechanical metastructure to enable controllable passive deformation. The metastructures and bone braces are designed to affix the femur bone to the tibiotarsus, whilst still permitting kinematic movement between the tibiotarsus and the tarsometatarsus of the crow skeleton. The rearticulated hips function as a fulcrum between the upper and lower body parts, whilst concurrently enabling sagittal rotation of the crow skeleton about the hips. Static compression tests, finite element analyses, and in-situ tests conducted using Sundoli shows that the deformation behaviours of metastructures with and without supports are acutely sensitive to the angle of the tarsometatarsus relative to both the ground and the loading direction, highlighting the importance of designing the metastructure holistically and with consideration to the entire skeletal structure. At different loads and angles, the metastructures exhibit variable stiffnesses over their full deformational ranges, demonstrating their effectiveness in protecting the brittle biological bones. Using a metastructure as a mechanism for passive joint rearticulation enables Sundoli an ability to support a payload 8.7 times its body weight without lateral support (an 870% payload ratio) and 14 times its body weight with lateral support (a 1400% payload ratio). This payload capacity is achievable through the full range of its upper body movement in the sagittal plane throughout the full range of its upper body movement in the sagittal plane.

## I. Introduction

Biological body parts can be roboticised [1] either as cybernetic organisms, where electrical stimuli are used to control physical movements, or as necrobots (necro-robots), where parts of the dead animal are bionically engineered to create a robot. Each of these areas conjoins the unique properties of natural materials with precision engineering and advanced methods of control. Insects, such as spiders, beetles, and cockroaches, are some of the animals that have been used to create hybridised biological robots. Tsvetkov and Alam for example [2], mechatronically optimised a rhinocerous beetle, enabling it to bear significant weight and achieving at its measured maximum, a payload ratio of 6847% (68.47 times its own weight). Yap et. al. [3] designed a gripping/grasping actuator from deceased spiders, which was able to lift 130 times its mass. Jørgensen et. al. [4] developed reinforcement mechanisms to enable the rearticulation of a canine skull for use as a robotic end effector. Conformable electrical interfaces and Venus fly traps were used by Li et. al. [5] to develop actuators, and Sanchez et. al. [6], Shoji et. al. [7], and Rasakatla et. al. [8] designed cyborg cockroaches with a functional gait that could have potential in search and rescue operations. Liu et. al. [9] focused on developing the quality of communication between the cyborg and the user, which can have knock-on benefits to the accuracy of physical movements such as those discussed by Nguyen et. al. [10], who engineered both posterior-to-anterior as well as lateral motion into a cyborg cockroach to optimise its global manoeuvrability in potential search-and-rescue type missions, for which cyborg beetles are also under development [11], as are those that have been engineered for flight [12].

Necrobotics to date, is a less researched area than the area of cybernetic insects. Yet, necrobots have the potential for utility in several application areas, as animal body parts are pre-formed structures suitable for application in robotic design. Robotic components sourced from deceased animals are an effective example of environemtnally sustainable robotic design, as the biological component is entirely biodegradeable. The integration of machine technology with nature is made easier through advanced manufacturing methods such as additive manufacturing (AM). Fused deposition modelling (FDM) is one of the many AM methods available, and polymers such as polylactate (PLA) can be used to produce complex parts with a reasonable level of strength [13] compared to materials like bone [14], and at a reasonable level of accuracy. AM offers reductions in production cost due to the democratisation of commercially available FDM 3D printers as well as material waste, as there is no post processing or other material needed, thus promoting a circular economy [15]. Such manufacturing methods compliment the environmental ethos of necrobotics, which is to use available resources to both democratise and make accessible, the engineering of functional robots.

A significant consideration in the design of necrobots is limb rearticulation and joint reconstruction. Since the original soft tissues such as muscle and cartilage can no longer be used in necrobots, mechanical joints need to be designed and integrated within the skeletal system. Metastructures hence have the potential for use as a rearticulation mechanisms since they are designed to enable a desired global property or behaviour. There is potential in the utility of these to simplify the control of robotic structures and their actuation. There is already a large number of examples where metastructures have been succesfully used in robotics. Hu et. al. [17] for example, designed an origami-inspired metastructure-based robotic arm for a crawling robot, while computable motion has been shown to be possible in an auxetic metamaterial robot [18]. Asymmetric and magnetic metamaterials can also enable crawling and swimming locomotions, [19] while crawling soft robots have been designed based on auxetic buckling-driven metamaterials [20]. The application of metastructure-robotic interfaces can be translated to the naissant field of necrobotics, providing a valuable solution to rearticulation controllable rearticulation, actuation, and joint reconstruction.

In this paper, we design discontinuous geared mechanical metastructures as applied to one pair of leg joints in a crow endoskeleton. We research the structure with and without the presence of the crow endoskeletal structure with respect to deformations controlled by stiffness, which is designed to change as a function of imposed loading and loading angle.

## II. Methods

### A. Design and Manufacture

#### 1) General details

Sundoli, shown in Figure 1(a), is a necrobot manufactured from a crow skeleton. It includes a passive articulation mechanism between its tibiotarsus and tarsometatarsus. This mechanism is designed to exhibit different stiffnesses and deformations as a function of loading and loading angle, and is enabled using geared mechanical metastructures. The metastructure acts as both a mechanism for passive joint articulation and energy absorption, allowing the necrobot to move its body nonlinearly whilst retaining a payload at different angles of upper body orientation. Each gear pair (upper and lower) is located on the tibiotarsus and tarsometatarsus, respectively, on each of the right and left legs. A truss within the metastructure forms the loading connection between the two bones. The femur is connected directly to the tibiotarsus via a web between the braces of the two bones. The braces, as well as the hip joint, are manufactured from FDM prints using polylactic acid (PLA) that enables rapid manufacture, considering the complex geometry of the metastructure. A servo motor (SM-S2309S, micro analogue servo with 4 plastic gears and 1 metal gear) was attached to the legs to control the angle of the body with respect to the legs by pushing against the ventral surface of the spine. The leg (femur, tibiotarsus, and tarsometatarsus) and the metastructure permits one degree of freedom (DoF) of motion. This is a rotational motion that changes the relative angle between the tibiotarsus and the tarsometatarsus *α*, while the rest of the body rotates around the hip-femur joint (i.e. the body angle) *θ*. We considered two design iterations (Iteration I and II), the second of which was an updated design based on test results from the first.

**Fig. 1.**
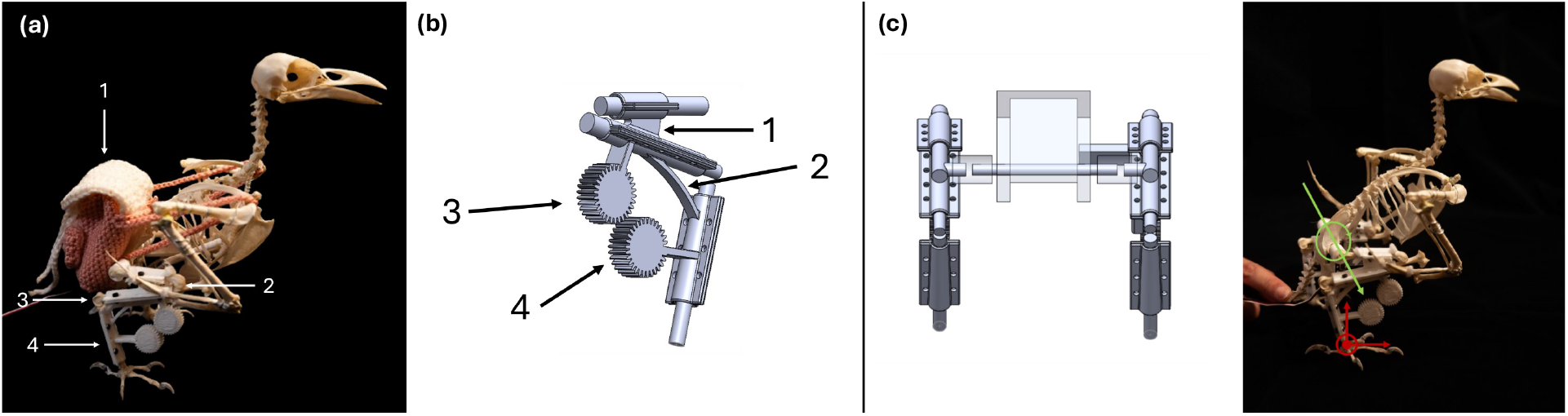
(a) Sundoli with backpack (1) holds weights, (2) is the femur, (3) tibiotarsus, and (4) tarsometatarsus, (b) CAD model of femur, tibiotarsus, and tarsometatarsus and its braces. (1) is web between the femur and tibiotarsus, (2) is the connection truss, (3) is the top gear, and (4) is the bottom gear, and (c) hip structure for hip articulation (left) and origin and axis (red), as well as rotation vector for body rotation in green (right).

#### 2) Metastructure DesignIteration I

The passively articulating joint is designed to accommodate geared mechanical metastructures and curved truss supports. These are connected to both the tibiotarsus and tarsometatarsus as shown in Figure 1(b). The curved truss supports are dimensioned to bear the full weight of the crow skeleton whilst concurrently providing a separation distance between the gears, such that there is no gear meshing until the gears are deformed to a specific load and angle. At a critical load (with respect to angle), the gears mesh and absorb energy from loading. The combination of the two structures gives the joint the potential to passively vary its stiffness while concurrently exhibiting an alternating range of motion. The gears were designed to be wide enough to enable meshing even if there are any offsets in loading or bone misalignment.

#### 3) Metastructure Design: Iteration II

After initial testing, we noted that there were several possible design improvements to Iteration I of the metastructure. The hip articulation mechanism was markedly unstable under load, leading to bone-to-bone contact. The inherent stiffness of the bones when in contact prevented the metastructure from being used to its full potential, and posed a further risk of damaging the bones with additional loading. Therefore, new structures were added to stabilise the hips, strengthen the spine, and potentialise a wider range of deformational motion in the metastructure. A cross-bar was attached to the two gear sets between the two legs to maintain the distance between them and only allow the indented in-plane motion, adding stability to the hips. The cross-bars were designed with an hourglass shape to minimise weight whilst maximising the surface area of attachment to the gears. Holes were placed within the gear trusses and connection trusses to decrease the stiffness, save weight, and encourage elastic load-deformation behaviour with variable stiffnesses. This second iteration of the metastructure is shown in lateral view in Figure 2(a), which also shows the position of the spine block, cross-bar, loading block, and the weight-saving holes (Figure 2(b)). A more detailed view of the second iteration of the structure is shown in Figure 3 from anterior (a) and posterior (b) views.

**Fig. 2.**
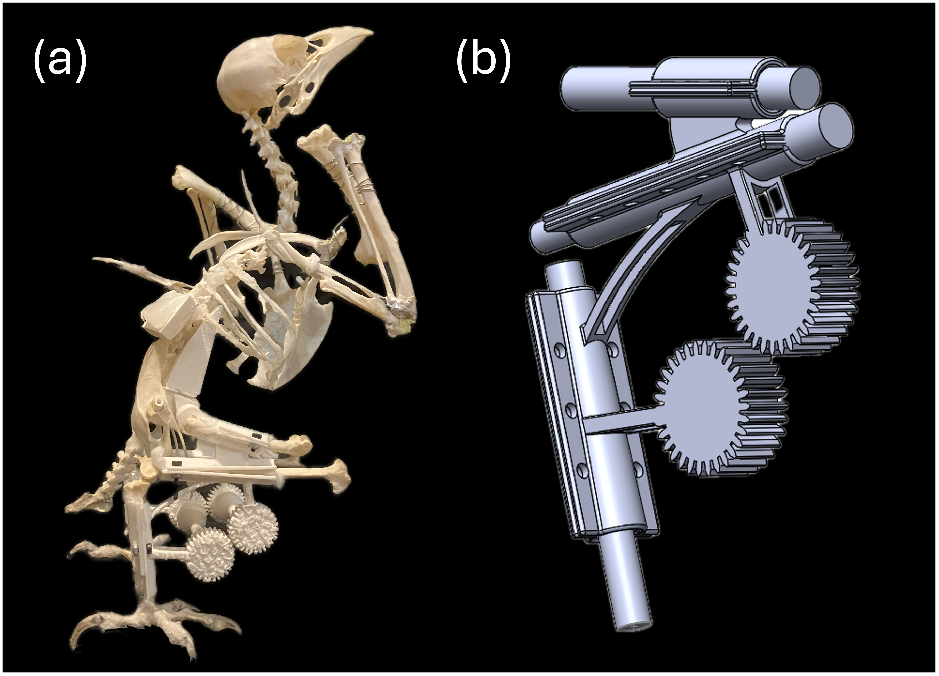
(a) Lateral-view of Sundoli with the Iteration II of the metastructure and (b) weight-saving holes in trusses.

**Fig. 3.**
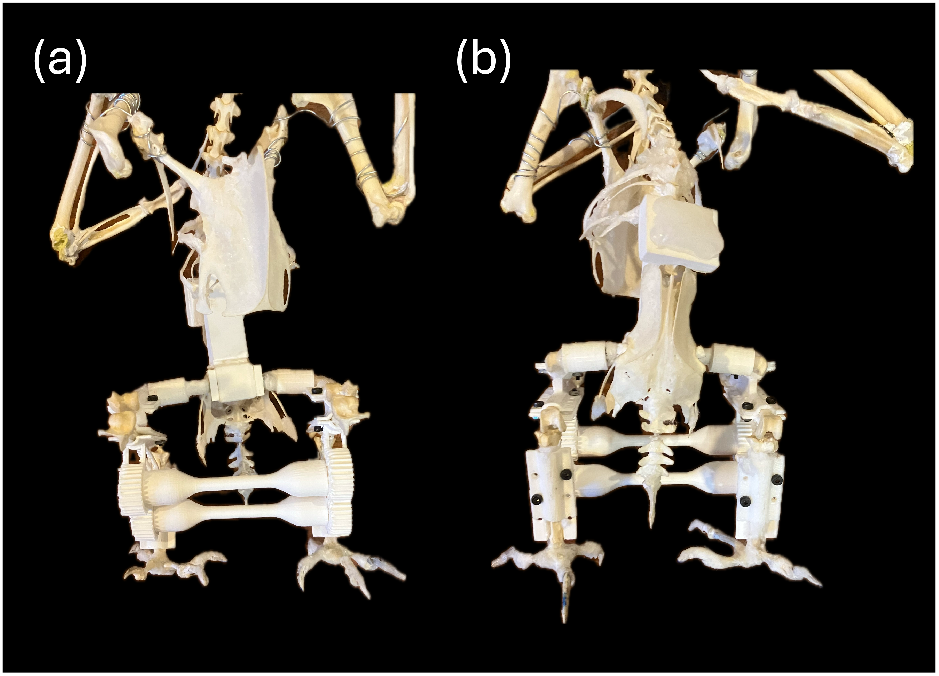
(a) Anterior view of Sundoli showing the cross-bars and spine block and (b) Posterior view of Sundoli showing the force adaptor on the spine.

#### 4) Articulation & Motion Design

The femur is secured to the tibiotarsus with a thick web between the braces of the metastructure, as shown by (1) in Figure 1(b), restricting its translation and rotation relative to the tibiotarsus. The tarsometatarsus is connected to the tibiotarsus through the truss connecting the two braces as shown by (2) in Figure 1(b) with (3) and (4) indicating the top and bottom gears respectively. Figure 1(c) displays the hip articulation mechanism in the crow, as well as the centre of rotation and the axis used. The centre of rotation of the body is shown in Figure 1(c) in green. The *x*-axis is towards the head of the crow, *y*-axis towards its beak, and *z* follows the right hand rule.

The braces cover the majority of the bone length between the top and bottom brace and are secured using M2 bolts. Cotton pads were placed between the braces and the bones to (a) dissipate mechanical energy and thus delocalise stress concentrations on the bone, and (b) account for variance in bone shaft diameter. The servo motor was used to rotate the body of the crow, thereby controlling the DoF of the body with respect to the leg. The servo is in turn is attached to the tibiotarsus brace in order to ensure that the reaction moment from body rotation is transmitted through the fixed parts of the structure. A lever was attached to the servo moment arm to improve the rotational capacity and stability of the crow body. This lever was designed to interface with the ventral face of the vertebrae between the foramen and parapophysis.

A computational scheme of the mechanism designed for hip articulation is shown in Figure 1(c). A hollow cylinder was fixed within the cavity of both of the hip joints with a steel wire running through them, each of which was connected to a hip cap. The cylinder, 3D printed in PLA, prevents the steel wires from wearing through the hip bones during use. While the hip cap fits over the cylinder, it is permanently adjoined to the wire, which enables rotation due to the servo. The opposite end of the hip cap was also permanently affixed to the femur.

### B. Static Compressive Loading

#### 1) Isolated Metastructure Testing

Compressive loads were applied to 8 samples (*n* = 8) to map the force-deformation response of the geared metastructure through its full deformation cycle. The bones were replaced with steel rods of similar diameter to isolate the effect of the metastructure in these tests. Steel rods were glued to an acrylic plate to reproduce a fixed boundary condition of the in-situ test as closely as possible. Two sloped 3D-prints were connected to the bottom of the acrylic plate to angle the bones in a similar manner to the angle observed in Sundoli. The metastructure was isolated in this test to more effectively assess its discrete behaviour under compressive loads, which were applied at 10 mm/min to the top surface of the tibiotarsus brace in an Instron 3369.

### C. Finite Element Analysis

The isolated metastructure was analysed using the finite element analysis (FEA) method under static loading conditions and using an implicit solver. A high material print density (i.e. the print infill density) was used to assume material isotropy and therefore simplified the simulation. The material was modelled as an elastoplastic solid with the following properties taken from Letcher et. al. [21]: density, *ρ* = 1250 kgm^−3^, Young’s modulus, *E* = 3.6 GPa, Poisson’s ratio, *ν* = 0.4, yield stress, *σ*_*y*_ = 64.03 MPa. The bottom edge of the tarsometatarsus was fixed in all DoF and a compressive displacement applied to the top of the tibiotarsus brace at a distance that was four times greater than the displacement applied during experimental testing. This was done in part, to test the performance limits of each metastructure. A two-way (both contact surfaces are calculated as master and slave in one timestep) frictionless contact was applied between the upper and lower gears alongside tied contact between the braces to one another, and to the bones to restrict displacement between assembled structures. The element used for the rigid parts representing the bone was a constant stress solid element, used for efficiency and accuracy. The braces were modelled with 1 point tetrahedrons for efficiency and enhanced geometry representation. Hourglass control was added to the model to account for the reduced integration points.

### D. In-situ Compression Testing of Metastructures

Static compression testing (Figure 4) was then conducted to assess the behaviour of the metastructure when incorporated within Sundoli’s skeleton. Load was imparted directly to a transmission block (25 by 20 mm) affixed using hot glue to Sundoli’s backbone over the thirteen to fifteenth vertebra to reduce stress concentrations and to distribute the load more evenly to the skeleton. The skeleton incorporating the metastructure was loaded and unloaded statically 8 times (*n* = 8 to determine whether the metastructure remained in an elastic state during its loading cycle. The strain rate was reduced to 5 mm/min to minimise chances of the bones breaking. A curved PLA wedge was hot-glued to the ventral face of the spine from twelfth vertebra to parts of the hip cavity to preserve the vertebral structure under loading, since the wire articulating the spine is easily deformable and this may affect the efficiency of load transmission to the metastructures. The body angle was kept constant by wire suspension.

**Fig. 4.**
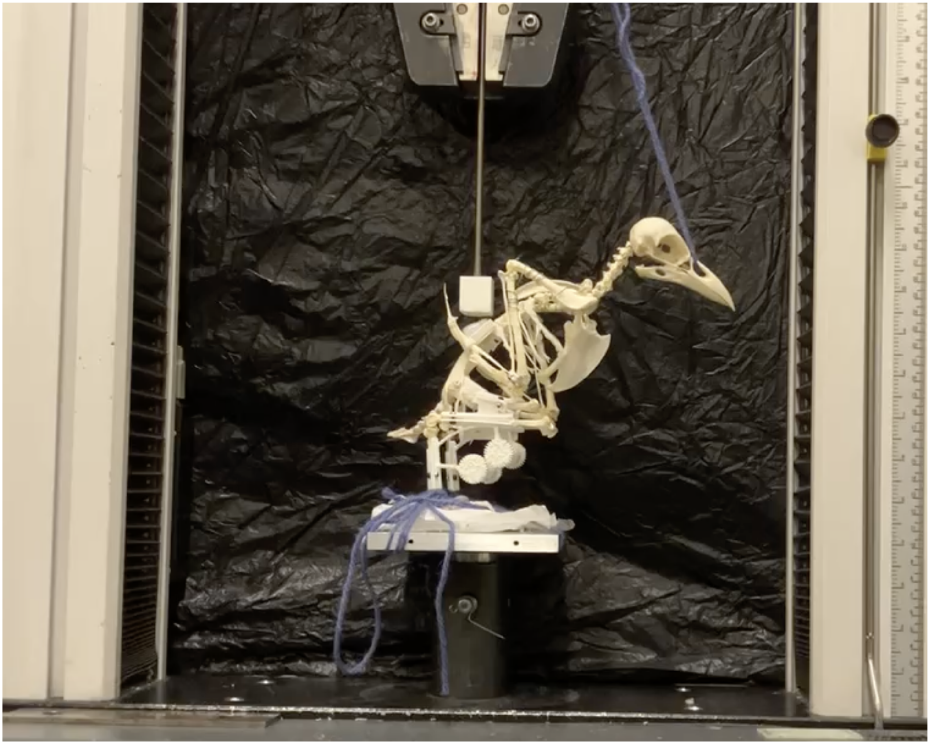
In-situ static compression testing set-up.

### E. In-situ Effects of Dynamic Compression Angle on Metastructure Kinematics

Steel ball bearings weighing 320 g were placed into Sundoli’s backpack (cf. Figure 1). The effect of backpack loading on the kinematics of the metastructure was filmed and tracked using Tracker, an open software created by Douglas Brown et al. based on the Open Source Physics (OSP) Java framework. Sundoli’s body angle, *θ*, was then varied in Iteration I using a servo motor held by the tibiotarsus brace and attached to a lever that would control the body motion of the skeleton. The servo was connected to an Arduino Uno which was used to input an angle of rotation, *θ*, to the servo (and hence the bird skeleton via the connected lever). These tests were conducted to monitor the influence of angular compression on the kinematics of the metastructures. In Iteration II, the cross-bars between the metastructure restricted the utility of the internal servo and thus motion was controlled externally via a wire inserted into Sundoli’s nasal cavity.

## III. Results and Discussion

### A. Isolated Metastructure Testing

The metastructure was noted to be non-linearly elastic when loaded up until failure. We simplified the different elastic behaviours using four linear constant from the force-deformation curves (A-D) as shown in Figure 5. The behaviour observed in (A) is an initial linear elastic stiffness arising from the elastic bending of the connection truss between the tibitotarsus and the tarsometatarsus. In (B) The upper gear moves towards the lower gear and meshes, causing the bottom gear truss and connection truss to bend further. The stiffness observed in region (C) arises through contact between connection truss and tibiotarsus brace. In (D) there is a linear strain-stiffening observed prior to failure, which is due to the storage of additional mechanical energy in the metastructure through a combination of meshed tooth shearing, truss bending and contact between the tarsometatarsus brace and the loading plate. The different stiffnesses in each of the regions (A-D) are provided Table I. There is significant variation in the values of stiffness in region D due to the nature of the deformation during failure and manufacturing method. Because the additive manufacturing method leads to an anisotropy within the structure, the failure mode of the eight samples are statistically different to one another. While the coefficient of variation in region C is high, this is due to the relatively lower stiffness value.

**TABLE I.** Stiffness calculations from isolated metastructure static compression testing.

| Coefficient | A | B | C | D |
| --- | --- | --- | --- | --- |
| Stiffness, $k = \frac{\Delta F}{\Delta \delta_c}$ [N/mm] | 3.68 | 6.9 | 2.2 | 22.85 |
| Standard Deviation [N/mm] | 1.19 | 0.97 | 1.72 | 12.2 |
| Standard Error [N/mm] | 0.42 | 0.34 | 0.61 | 4.31 |
| Coefficient of Variation [%] | 32.3 | 14.3 | 78.4 | 53.42 |

**Fig. 5.**
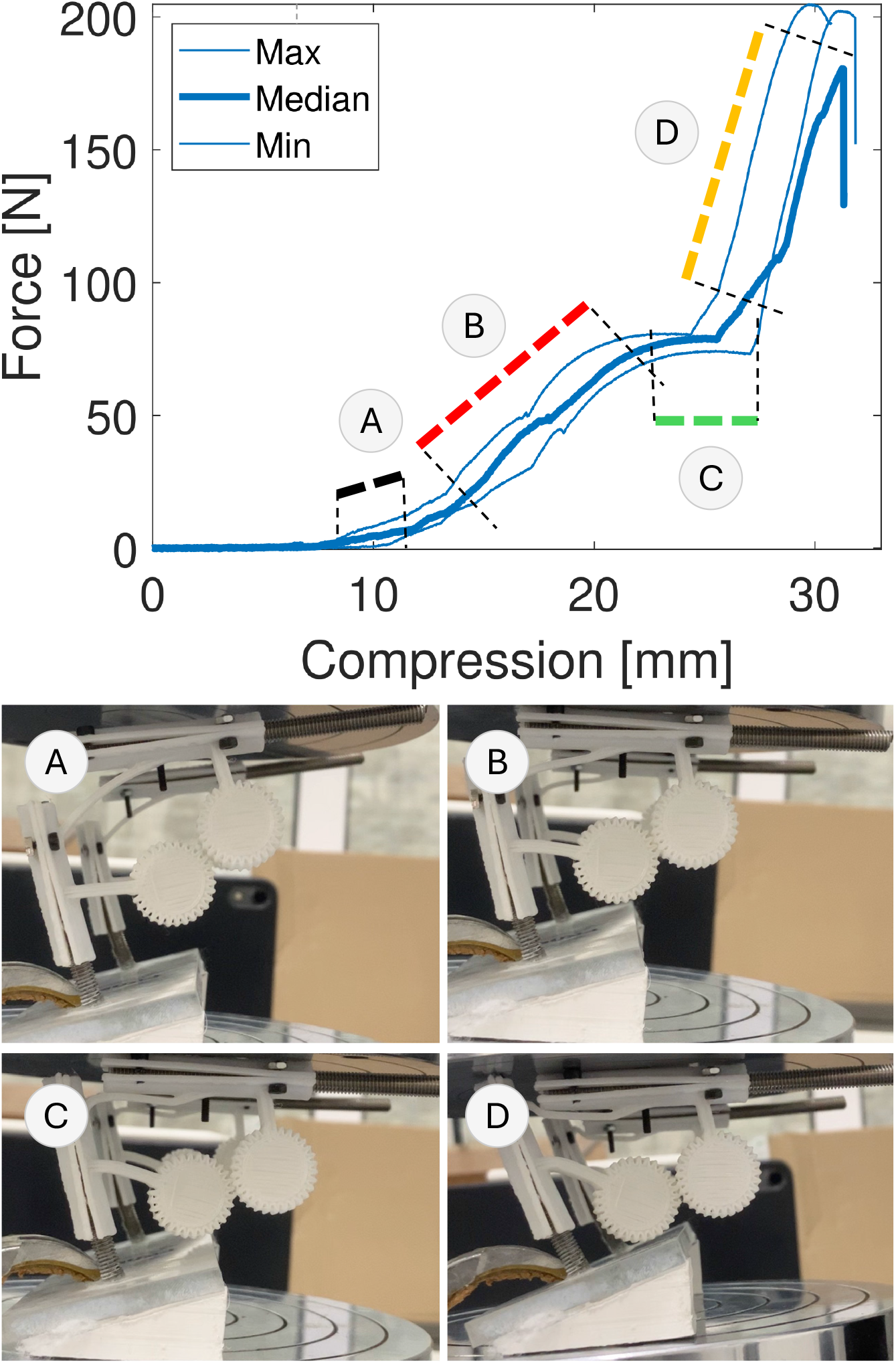
Force-displacement curve (force-compression curve) from static testing. Deformation sequence from top left going clockwise.

#### 1) Kinematics

Figure 6(a) shows the kinematic motion of the gears through a complete compressive loading cycle. The upper gear travels dorso-ventrally with a slight anterior-posterior curvature due to a direct loading of the tarsometatarsus and the resultant angular displacement of this segment of the metastructure. The truss connecting the braces bends, allowing the upper and lower gears to mesh, which in turn enables lower gear truss flexure and a resultant tighter curvature in the kinematic motion is observed for the lower gear. This bending occurs because the gear cannot rotate and meshing generates a moment on the gear truss.

**Fig. 6.**
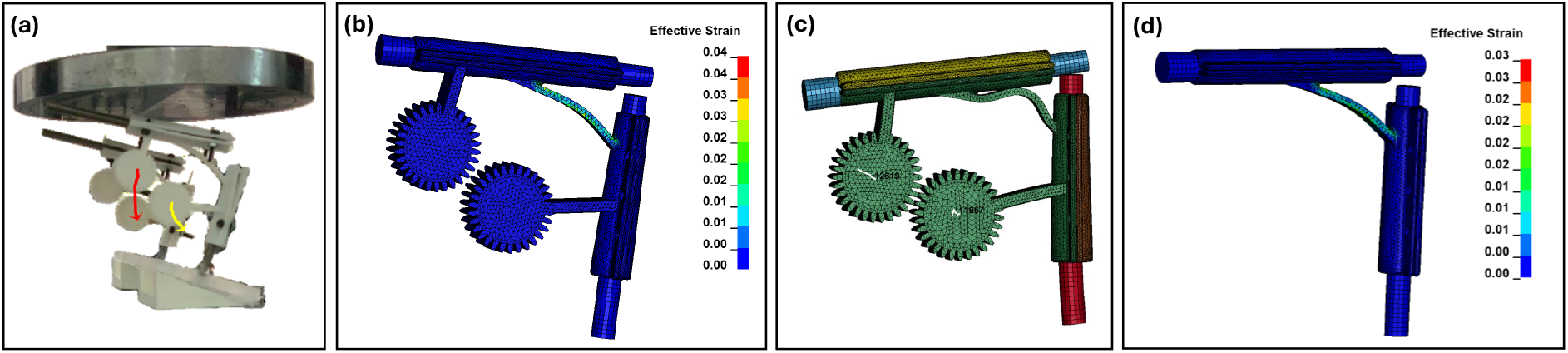
Metastructure Iteration I: (a) *x* − *y* motion of gears during experiment superimposed to gear location at start (b) strain of leg and metastructure as computed by FEA (c) *x* − *y* motion of gears from FEA and (d) strain of leg and metastructure in FEA without gears.

#### 2) Energy Absorption

Table II provides values for the energy absorbed, *W*, by the metastructure within each of the regions (A-D) calculated as *W* = ∫*F*d*δ*_*c*_, where *F* is force and *δ*_*c*_ is compressive displacement. The highest energy absorbed is in region (D), where the gears are meshed and the metastructure strain stiffens to failure. During testing, we observed that the meshing of the gears delayed the onset of failure in the connection truss by redirecting the deformation (and hence force) away from the axis of the machine crosshead. The metastructure fails at the interface between the connection truss and the tarsometatarsus brace. This was expected since it was the structure that endured the highest visible deformation during the test. The highest variability can be noticed in the energy absorbed under stiffness region C because the start and end points of these regions differ most prominently due to torsional behaviour of the metastructure under higher loads.

**TABLE II.** Average Energy Absorbed from isolated metastructure static compression testing

| Value | A | B | C | D |
| --- | --- | --- | --- | --- |
| Energy [J] | 0.3 | 0.262 | 0.169 | 0.582 |
| Standard Deviation [J] | 0.07 | 0.124 | 0.35 | 0.303 |
| Standard Error [J] | 0.025 | 0.044 | 0.124 | 0.107 |
| Coefficient of Variation [%] | 23.48 | 47.2 | 207.2 | 52.2 |

### B. FEA

A 1 mm sized element was used following mesh convergence between 0.4-2 mm as it yielded consistent energy and displacement results with the smaller element sizes. The simulated behaviour was similar to that observed in the static test, though minor differences were noted due to differences in the loading direction (more tangential to the connecting truss to the tibiotarsus), boundary condition (fixed translation on the underside of the tarsometatarsus), and material anisotropy (due to 3D printing). The connection truss buckles rather than bends in Iteration I of the metastructure, as was also observed in the experimental test, and the displacement of the gear shows reduced *y*-direction motion. The deformation and force is nevertheless similar in magnitude. Figure 6(b) reveals significant straining on the connecting truss during deformation when the gears are meshed in metastructure Iteration I. This is partly due to the higher displacement applied to the model compared to the experiments. There is no shearing of the gear teeth or the gear trusses at this stage, and further deformation is limited by the contact between the tibiotarsus and tarsometatarsus, which is more representative of the in-situ testing configuration. When comparing the strain experienced in metastructure Iteration I against the added cutouts in the trusses in metastructure Iteration II, we find that cutting out material from the truss segment in metastructure Iteration II results in an increase in strain by 0.05 in the trusses, comparing Figure 7 against Figure 6(b). The simulations indicate that in both iterations, the connecting truss is the most deformable component of the metastructure, and may be the location from where failure initiates, should the structure fail plastically.

**Fig. 7.**
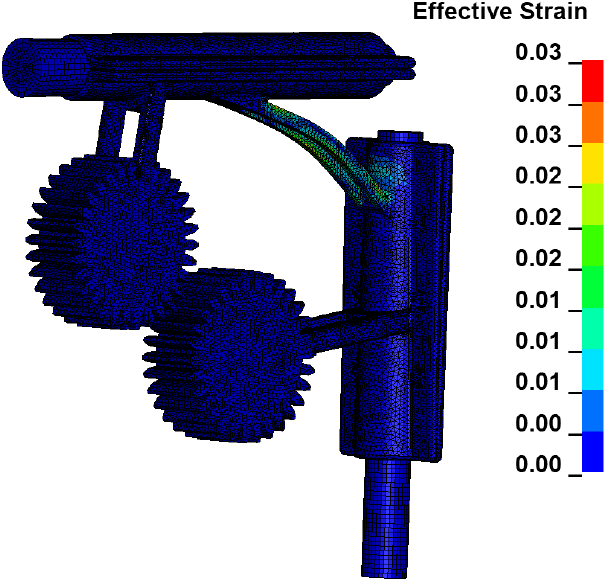
Metastructure Iteration II: strain of leg and metastructure as computed by FEA.

#### 1) Kinematics

The trajectory of the gears are shown in Figure 6(c). As there are no manufacturing imperfections, the gears mesh perfectly and rotate, causing the bottom gear to take a dorsal then ventral trajectory. There is significantly less *y*-motion as the loading direction is different to that of the static test.

#### 2) Energy Absorption

The same structure was simulated without the gears to analyse the effectiveness of the gears in absorbing mechanical energy. Figure 6(d) shows the strain without the gears at the same time step as Figure 6(b), and while the deformation shapes are similar, the maximum force value is at least two orders of magnitude lower. The strain value without gear in the connection truss is also 2% higher. The gears therefore are evidenced to play a significant role in energy absorption.

### C. In-situ Static Compression Testing of Sundoli using Metastructure Iterations I and II

Both metastructure iterations were tested attached to the skeleton. The crow skeleton was loaded from on its spine, enabling for passive deformation of the metastructures. Figure 8(a) shows the minimum, maximum, and the median curves obtained from the in-situ static compression test of metastructure Iteration I (where number of repeated test *n* = 8). Here, all specimens consistently show lower initial stiffnesses, A, (average of 0.68 N/mm) where gears were incompletely meshed and the entire skeletal structure was not locked in place. After some loading, the average secondary stiffness, B, increased to 1.74 N/mm, due to both the complete meshing of the gears, as well as interactions between the tibiotarsus and tarsometatarsus, and other skeletal regions. Figure 8 (b) shows the median, minimum, and maximum force-displacement curve from the in-situ compression testing of Iteration II. Contrarily Iteration I, the stiffnesses decrease from A to B (from on average 1.91 to 0.59 N/mm, where *n* = 8). The initial stiffness increase is due to gear meshing. However, due to the added stability of the cross-bar between the knees, less energy is used in the skeletal segment inter-locking, allowing more mechanical energy to be absorbed by the metastructure. Because of the weight-saving cut outs, it is likely that slightly less energy would be absorbed through deformation of material than in Iteration I. Nevertheless, since the first iteration is unstable, more deformational energy is passed to the skeleton of Sundoli and the metstructure is not used to its fullest potential. The results are summarised in Table III. The energy absorbed by Sundoli using each of the metastructure Iterations I and II, is shown in Table IV.

**TABLE III.** Stiffness calculations from in-situ static compression testing of both Iterations I and II of the metastructure.

| Iteration I | A | B |
| --- | --- | --- |
| Stiffness, $k$ [N/mm] | 0.68 | 1.74 |
| Standard Deviation [N/mm] | 0.17 | 0.91 |
| Standard Error [N/mm] | 0.06 | 0.32 |
| Coefficient of Variation [%] | 25.2 | 52.3 |
| Iteration II | A | B |
| Stiffness, $k$ [N/mm] | 1.92 | 0.59 |
| Standard Deviation [N/mm] | 0.63 | 0.47 |
| Standard Error [N/mm] | 0.22 | 0.17 |
| Coefficient of Variation [%] | 32.7 | 79.7 |

**TABLE IV.** Energy absorbed by metastructure and skeletal system during in-situ compression testing for both Iterations I and II of the metastructure.

| Iteration I | Value |
| --- | --- |
| Energy [J] | 0.029 |
| Standard Deviation [J] | 0.013 |
| Standard Error [J] | 0.0046 |
| Coefficient of Variation [%] | 44.9 |
| Iteration II | Value |
| Energy [J] | 0.014 |
| Standard Deviation [J] | 0.0033 |
| Standard Error [J] | 0.0012 |
| Coefficient of Variation [%] | 23.9 |

**Fig. 8.**
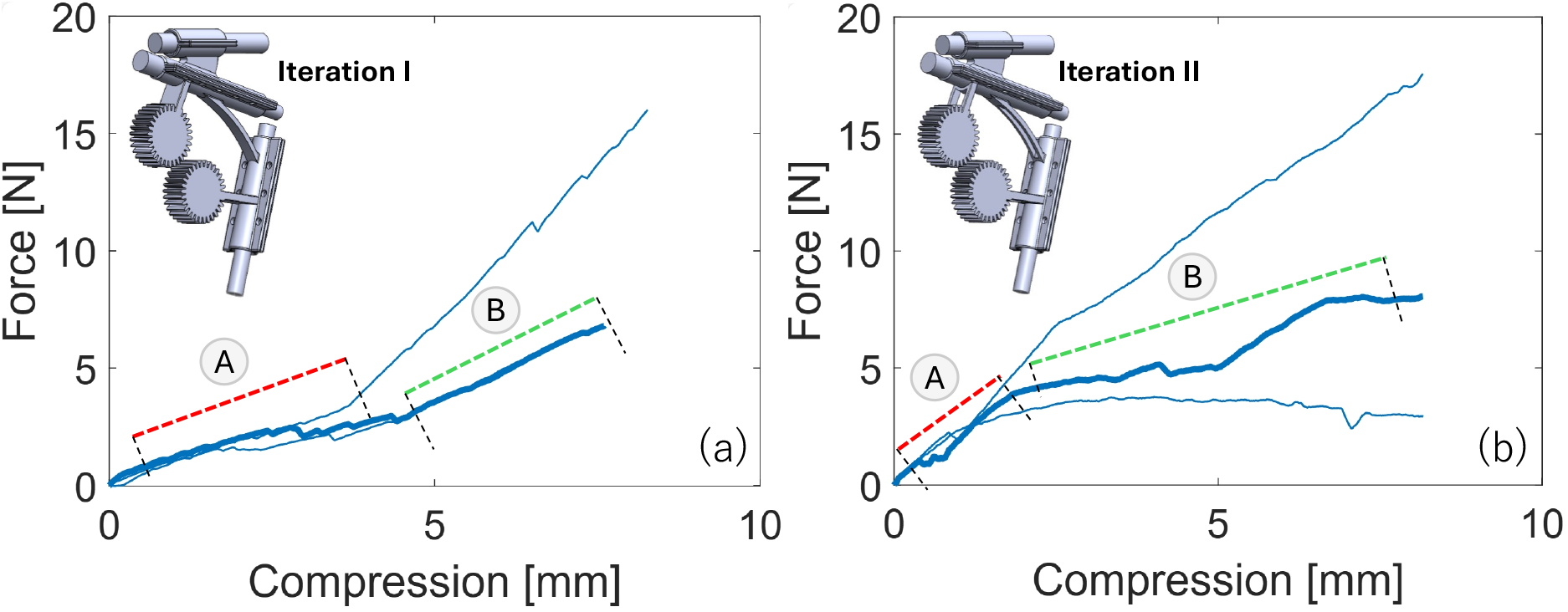
Force-displacement comparison between Iteration I (left) and II (right) under in-situ static compression.

### D. In-situ Angular Loading of Sundoli using Metastructure Iterations I and II

The metastructure and the hip articulation mechanism was assembled with the skeleton and loaded at different angles to test the metastructure in-situ. A weight was added to the back toes to provide additional stability and ensure that Sundoli would not topple over at lower *θ*. 320 g of weights were added to the backpack for Iteration I, which was placed in the middle of Sundoli’s body. The angle was varied from 60° to 20° using the servo-lever system, rotating from an almost upright position, to full anterior flexion.

#### 1) Kinematics

Under loading, the hips of Sundoli shift ventrally and loads are imposed upon the femur. As the angle is varied from 60°to 20°, there is a change in load magnitude and direction (force vectors) that are experienced by the metastructure. The loading becomes progressively more distributed across the back of Sundoli, shifting its load application location higher up the spine as the body becomes more horizontal. This results in a higher moment in the overall body. The force vector components along the spine also become smaller, enabling weight distribution to the metastructure, resulting in the meshing of the gears. The motion of the gears during compression is shown in Figure 9 for each of Iterations I (V1) and II (V2) as the angle is changed from 60°to 20°. There is less displacement in both *x* and *y* in Iteration II compared to Iteration I. This is due to increased lateral stability in the legs and hips, which is primarily due to the presence of the cross-bars (cf. Figure 3). By stabilising the lower body in this way, loading applied to Sundoli can be directed to the metastructure as the skeletal structure moves less. Displacement is observed to increase rapidly when the body angle reaches around 40°since the loading path approaches collinearity with the angle of the tarsometatarsus. With the added cross-bar and hip structure in Iteration II, Sundoli was stable enough to take 194 g more weight (514 g) than Iteration I.

**Fig. 9.**
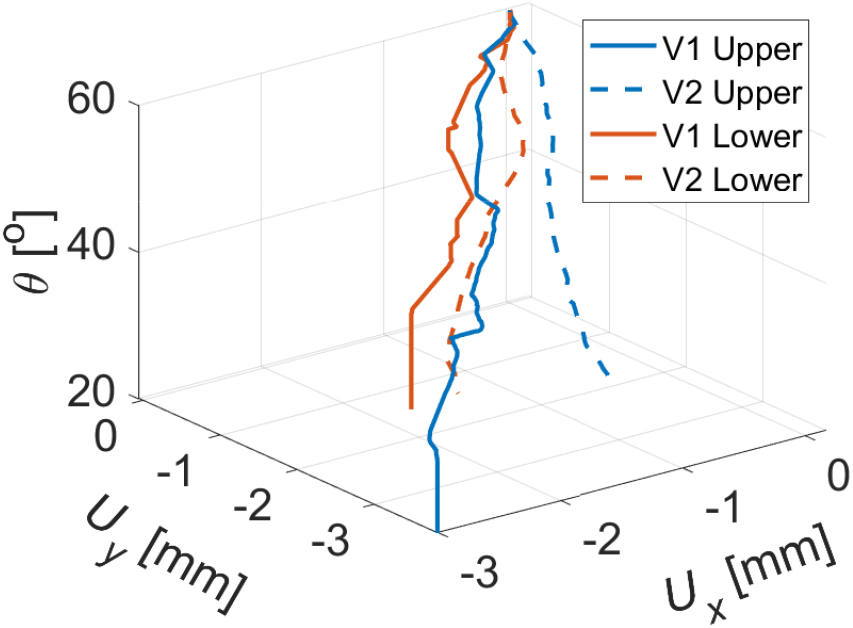
Gear motion in in-situ testing with weights.

#### 2) Energy Absorption

The metastructure Iterations I and II give the necrobot the ability to support and move with 8.7 times and 14.0 times its body weight (i.e. a payload ratio of 870% and 1400% respectively with the skeleton itself weighing 36.8 g), without experiencing any damage to the legs or toes. While the majority of the weight was supported through the metastructure, some would have been dissipated through the deformation of the hip articulation mechanism. If the hip articulation mechanism was more rigid, the metastructure would be able to absorb higher levels of mechanical energy, though there is a fine balance to the design of this as it may also lead to damage in the hip bones. Further testing is necessary with different materials and geometries, to ascertain the true load bearing limit states of Sundoli. When cyclically ramping the system, we also noted that there was no change in load-deformation behaviour.

## IV. Conclusions

A multi-stiffness necrobot joint has been designed and manufactured using geared mechanical metastructures. Two metastructure design iterations were analysed through static testing, FEA, and incorporated into a crow endoskeleton for insitu testing under both static compression and dynamic angular loading. Static tests and FEA on the isolated metastructure provides insight into its mechanical properties, behaviours, and kinematics under loading, each of which is intimately related to the loading direction relative to the angle of the tarsometatarsus. The gears enable heightened levels of mechanical energy absorption and can be used to passively control variable stiffness in the metastructure as a function of deformation. In-situ testing evidences that both metastructure Iterations (I and II) are successful in absorbing mechanical energy while protecting the crow skeleton from damage.

We show that variable stiffness and deformation can be designed into a necrobot articulation structure, enabling (in this work) a payload ratio of over 870% for a metastructure excluding lateral stabilisation, and a payload ratio of 1400% for a metastructure with lateral stabilisation. The high interdependence of the kinematics of the gear with the overall behaviour of the metastructure, and hence the movement of the crow body, is recognised as being implicitly related to the change in loading direction relative to the angle of the tarsometatarsus, as this relationship controls the final gear trajectories. This work demonstrates a first-of-its-kind endoskeletal metastructure and, as evidenced, has the potential to enable necrobots to move freely while carrying high payload ratios.

## Acknowledgment

We would like to thank Mr. Colin Kessler for his help in taking pictures and videos of Sundoli. We would also like to thank Mr. Calum Melrose and Mr. Dan Coupe for their advice on making the static compression jig. We thank Dr Amer Syed for comments on an early version of this manuscript.

## Notes

### Competing Interest Statement

The authors have declared no competing interest.

